# Expression of soluble methane monooxygenase in *Escherichia coli* enables methane conversion

**DOI:** 10.1101/2021.08.05.455234

**Authors:** R. Kyle Bennett, Nyaradzo Dzvova, Michael Dillon, Stephanie Jones, Kelley Hestmark, Baolong Zhu, Noah Helman, Derek Greenfield, Elizabeth Clarke, Eleftherios T. Papoutsakis

**Affiliations:** Department of Chemical and Biomolecular Engineering, University of Delaware, Newark, DE, USA; The Delaware Biotechnology Institute, University of Delaware, Newark, DE, USA; Industrial Microbes, Alameda, CA, USA

## Abstract

Natural gas and biogas provide an opportunity to harness methane as an industrial feedstock. Bioconversion is a promising alternative to chemical catalysis, which requires extreme operating conditions and exhibits poor specificities. Though methanotrophs natively utilize methane, efforts have been focused on engineering platform organisms like *Escherichia coli* for synthetic methanotrophy. Here, a synthetic *E. coli* methanotroph was developed by engineering functional expression of the *Methylococcus capsulatus* soluble methane monooxygenase *in vivo* via expression of its cognate GroESL chaperone. Additional overexpression of *E. coli* GroESL further improved activity. Incorporation of an acetone formation pathway then enabled the conversion of methane to acetone *in vivo*, as validated via ^13^C tracing. This work provides the first reported demonstration of methane bioconversion to liquid chemicals in a synthetic methanotroph.

## Introduction

Abundant and recoverable reserves of natural gas, as well as increased biogas generation, has provided the economic and environmental opportunity to utilize methane as an industrial feedstock for liquid fuel and chemical production ^1^. Compared to chemical conversion of methane to liquid fuels and chemicals via catalysis, which require extreme operating conditions and exhibit poor specificities, biological conversion of methane is a promising alternative ^2^. Though native methanotrophs possess the ability to utilize methane, engineering synthetic methane utilization in established platform hosts like *Escherichia coli* is preferred industrially ^3^. Though many groups have established synthetic *E. coli* methylotrophs that possess the ability to utilize methanol ^4–7^, no one has yet established a synthetic *E. coli* methanotroph that possesses the ability to utilize methane *in vivo*. Recently, it was shown that the particulate methane monooxygenase (pMMO) retained enzymatic activity *in vitro* after expression in *E. coli* and purification ^8^. However, *in vivo* methane bioconversion in *E. coli* remains a significant challenge despite multiple efforts.

Here, we engineered the functional expression of the soluble methane monooxygenase (sMMO) to achieve the first step in the synthetic methanotrophic pathway: conversion of methane to methanol. Heterologous expression of the full and functional sMMO has been notoriously intractable ^9,10^. The six-subunit enzyme forms a dynamic complex that turns methane and molecular oxygen into methanol and water, using NADH as an electron donor (Fig. 1A). MmoXYZ form the hydroxylase, which contains a non-heme, di-iron catalytic center; it has never been functionally expressed in a non-methylotrophic host, and solubility has been identified as one potential cause ^9,11^ The reductase, MmoC, and a regulatory protein, MmoB, have been successfully expressed, and the last component, MmoD, inhibits sMMO activity when supplemented to lysates but its exact physiological role has yet to be determined ^10,12,13^.

**Figure 1.**
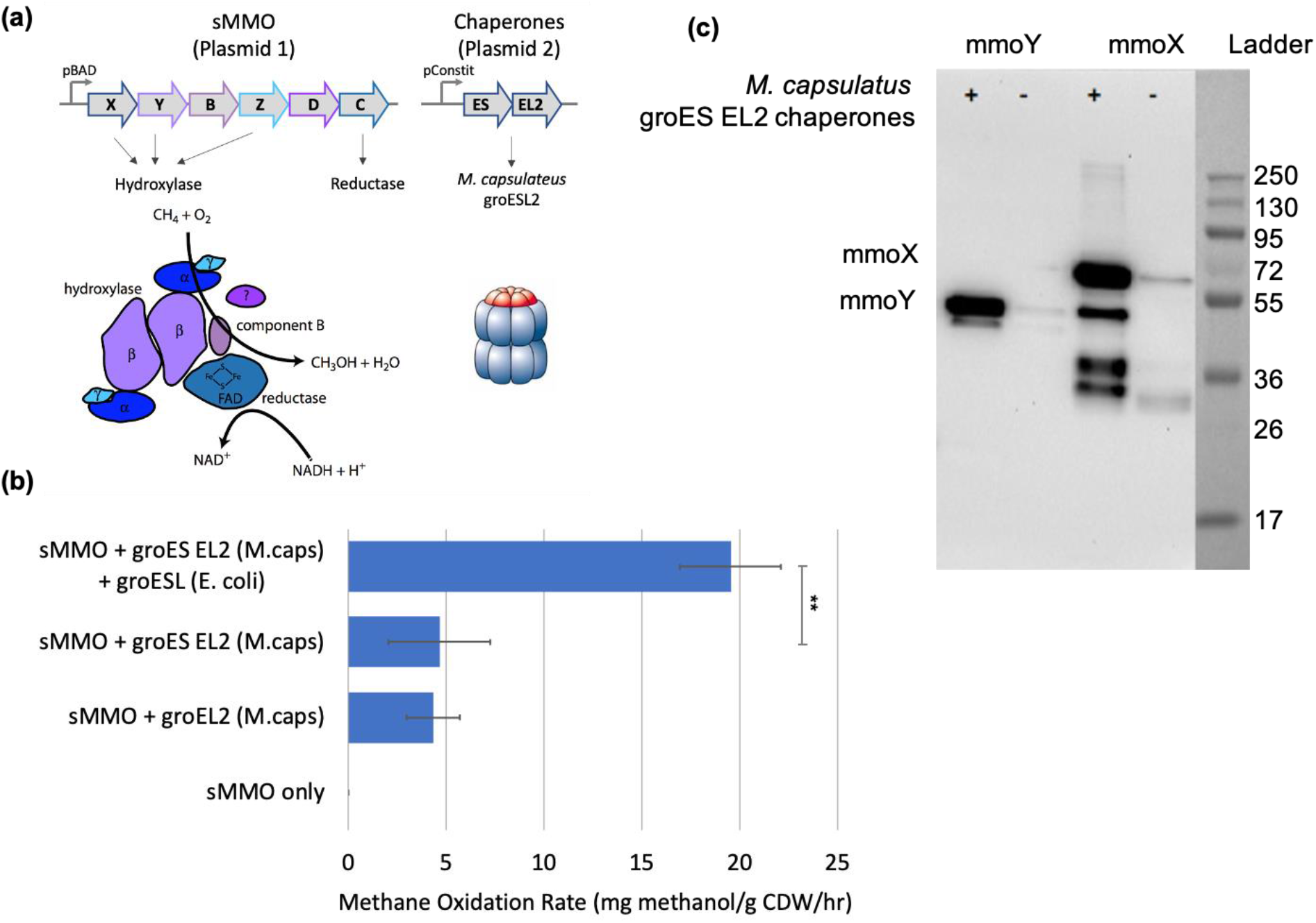
*In vivo* expression of an active soluble methane monooxygenase (sMMO) in *E. coli.* (a) Operon structure of the sMMO and folding chaperones. The sMMO converts methane and molecular oxygen into methanol and water, using NADH as an electron donor. Expression of folding chaperones, GroESL, is essential for MMO activity. GroEL (blue) is composed of two stacked seven-membered rings and interacts with the GroES lid (orange) to refold misfolded proteins. (b) *M. capsulatus* sMMO rate is chaperone dependent. (c) Expression of GroESL chaperones increases the amount of soluble MmoX (60.6 kDa) and MmoY (45.1 kDa). Error bars indicate standard error (n=5-6). ** p < 0.001. See text for more details.

Notably, there has been success in transferring active sMMO into heterologous methylotrophs. Lloyd, et al. demonstrated that a plasmid containing DNA from the *Ms. trichosporium* OB3b sMMO genomic locus conferred sMMO activity to *Mcy. parvus* OBBP and in *Mm. album* BG8, two organisms which only contain the particulate form of the MMO^14^. Though it is possible that additional genes in the recipient organisms contributed to sMMO activity, these results suggest that the complete set of genes necessary for sMMO expression and activity might be found in a single genomic locus. This and related approaches have enabled the mutagenesis and characterization of the hydroxylase component of the sMMO ^15,16^.

## Results

### Protein-folding chaperones enhance sMMO expression

Until now, the sMMO had not been actively expressed in *E. coli*. We suspected that the key to achieving functional expression was to test large sets of candidate sequences, including well-characterized MMO operons and protein-folding chaperones that might improve solubility. In natural methanotrophs, the genes encoding sMMO are most often found with genes encoding GroEL in the same operon, which suggested to us that this chaperone system in particular might be critical for proper sMMO expression. We synthesized a total of 17 candidate monooxygenases including methane monooxygenases, and monooxygenases for ethane, propane, alkene, and toluene (Table S1). We synthesized *groEL-ES* operons from the same organisms and designed a two-plasmid expression system that would allow us to test different combinations of monooxygenases and chaperones. We screened these using a sensitive fluorescent assay that relies on the promiscuity of the sMMO towards an alternate substrate: coumarin ^17^. We hypothesized that other functional soluble di-iron monooxygenases might also hydroxylate coumarin to umbelliferone, which can be measured via fluorescence.

We identified four active clones, demonstrating an active sMMO *in vivo*; characterization of these strains indicates that protein folding chaperones play a significant role in sMMO activity. Through sequencing analysis, these clones were shown to contain sMMO homologues from three different organisms, paired with either their cognate or non-cognate GroESL chaperones ^18^. Strains were validated for activity on methane by exposing cells in serum bottles to methane (supplementary methods) and quantifying methanol titers. We selected the highest-activity pair, the *M. capsulatus* (Bath) sMMO with its cognate GroESL chaperone, for further optimization. We hypothesized that the chaperones have a positive and necessary impact on the solubility of the sMMO complex. As shown in Fig. 1B, the *M. capsulatus* sMMO rate on methane depends on the expression of *M. capsulatus* GroESL2. Interestingly, expression of *M. capsulatus* GroEL2 alone is sufficient to generate a functional MMO, suggesting that the *M. capsualatus* GroEL2 forms a functional chaperone with *E. coli* GroES. Additional over-expression of *E. coli* GroESL further improved activity (Fig. 1B). Next, we sought to validate that chaperone expression indeed improves the solubility of sMMO. To do this, we expressed the entire operon with only one of the six subunits (*mmoXYBZCD)* his tagged, and the rest untagged. A total of 12 strains were constructed in order to combine each of the six plasmid constructs with or without a chaperone plasmid. As shown in Fig. 1C, the amount of soluble MmoX and MmoY increases dramatically when *M. capsulatus* GroESL2 is expressed. Additional bands are observed for his-tagged MmoX, suggesting some degradation products. Minimizing degradation could improve overall activity and is currently being explored. Other subunits are only moderately impacted (Fig. S1) and are soluble even without chaperone expression.

### Improving sMMO activity via directed evolution

We next improved sMMO activity via directed evolution. We generated a full site-saturation mutagenesis library on all six subunits of the sMMO operon (*mmoXYBZDC*) and the *groEL-2* gene. All 2,237 amino acids in these 7 polypeptides were mutated one at a time to all other 19 amino acids ^19^. A total of ~32,000 clones from this library were screened using the coumarin assay, representing an approximate coverage of ca. 50%. Many mutations were discovered that improved the rate of methane bioconversion above that of the wild-type enzyme; over 400 hits with improved activity were identified and validated ^20^. Of these, we picked 50 to recombine in a combination library, targeting an average incorporation rate of 5 mutations per library member. Coumarin data for these 50 mutants are shown in Figure 2A. This combination library was screened and the hits were validated for methane bioconversion (Figs. 2B–C). The best mutant from this library contained 5 mutations: MmoX^V23G,T356G^, MmoZ^R70E^, MmoB^Y139S^ and GroEL2^N409G^. It is currently unclear why these mutations improve methane oxidation, but due to the solubility issues, one hypothesis is that these mutations enhance the expression/solubility of the sMMO in the heterologous host. Finally, we moved this sMMO expression system into a set of *E. coli* strains that have mutations beneficial for folding and expressing challenging proteins (Table S2). The highest activity was found in OverExpress™ C43(DE3) (Figs. 2B–C), which exhibited a rate of 85 ± 2 mg methanol·gCDW^−1^·h^−1^. In comparison, natural methanotrophs expressing the sMMO have a reported rate of 100 to 1000 mg methanol·gCDW^−1^·h^−1^ ^21–24^.

**Figure 2.**
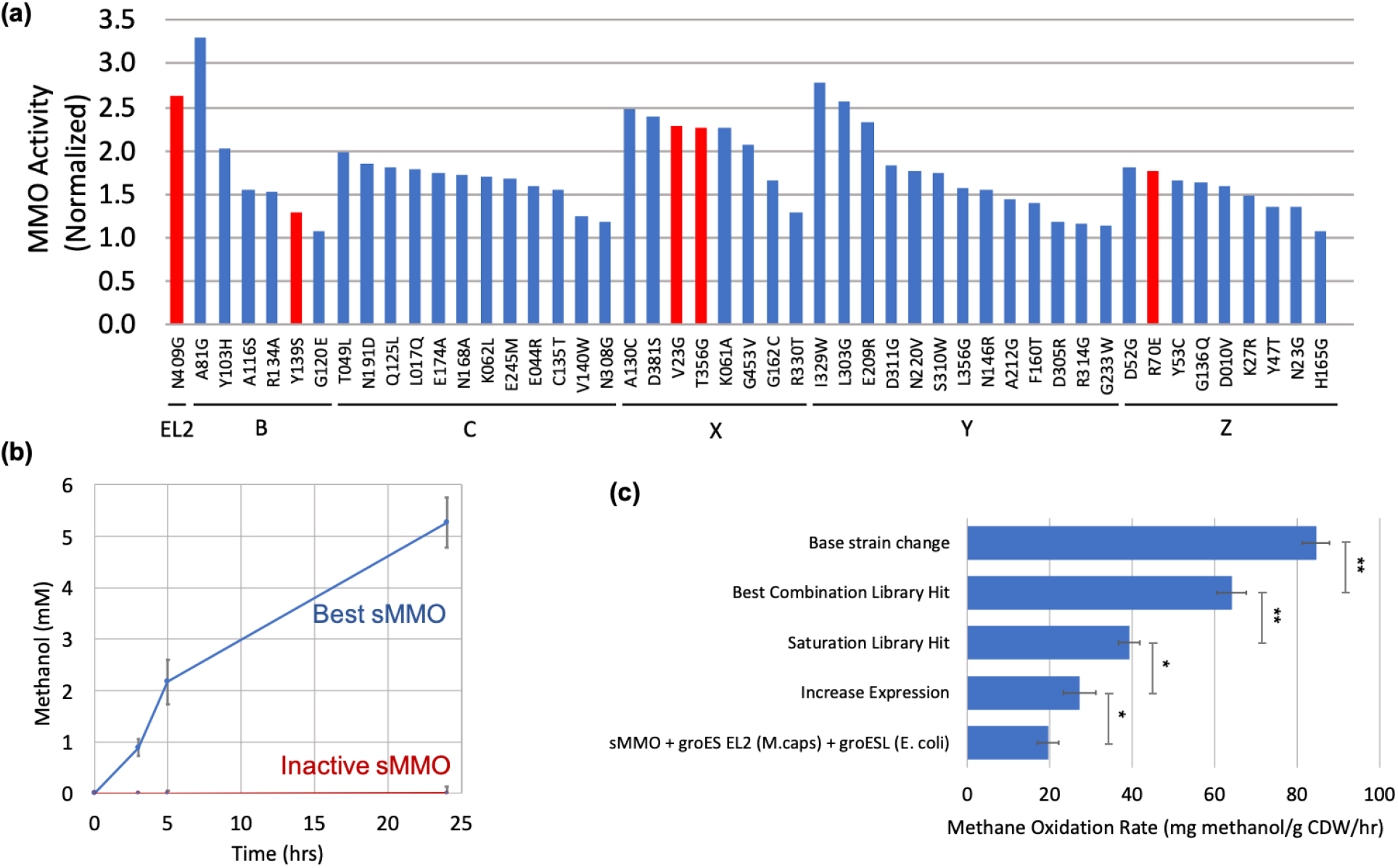
Improvement in sMMO rate via protein engineering. (a) Mutations that improve MMO MMO activity were identified by screening a site saturation library. Coumarin activity is normalized to a WT sMMO control. These 50 mutations were recombined in a combination library targeting an average of 5 mutations across the MMO operon and groEL2. Single point mutations found in the best combination library hit are shown in red. (b) *E. coli* expressing sMMO from *M. capsulatus* Bath converts methane to methanol. The initial rate between 0 and 3 hours is used to calculate a methane oxidation rate, which is plotted in panel c. A mutation in the catalytic site of MmoX (H246A) makes the enzyme inactive. (c) Improvements to the sMMO rate were found by combining the chaperones and the sMMO operon onto a single higher copy number plasmid (Fig. S2), and protein engineering via a full site-saturation library and combination library. An example of a single point mutation that increased activity found in the saturation library is shown here (MmoX^V23G^). The best combination hit was found to be MmoX^V23G,T356G^, MmoZ^R70E^, MmoB^Y139S^ and GroEL2^N409G^. Expression in a different *E. coli* strain (OverExpress) further improved the rate. Error bars indicate standard error (n=5-6). * p < 0.005, ** p < 0.001. See text for more details.

*E. coli* expression of the sMMO is a flexible and highly engineerable system for understanding and modifying this enzyme, that has advantages over the only other plasmid based heterologous expression system for this enzyme ^11^. Using this *E. coli* based system, we have shown the ability to change the substrate specificity for methane and ethane with a single mutation: MmoX^E240N^ (Table S3). By adding suitable origins to the sMMO plasmid, we demonstrated the sMMO exhibits *in vivo* activity in other, industrially relevant heterologous hosts as well, including *Pichia pastoris* (Fig. S3) ^18^. However, for the purposes of this study, we chose to focus on *E. coli* since there has been much more synthetic methylotrophy research performed in *E. coli* compared to other heterologous hosts.

### Conversion of methane to liquid chemicals in vivo

To examine if methane could be converted to liquid chemicals by a non-native methanotroph, we transformed the sMMO-expressing plasmid (pNH284) into a previously engineered *E. coli* strain (Δ*frmA*Δ*pgi* + pUD11) that co-utilizes methanol and glucose for acetone production ^25,26^. Expression of the functional sMMO in this strain realized the ability to co-utilize methane and glucose for acetone production (Fig. 3A). As demonstrated by the fermentation data with ^13^CH_4_ (and the N_2_ control) atmospheres, this engineered *E. coli* strain (Δ*frmA*Δ*pgi* + pUD11 + pNH284) can convert methane and glucose to acetone (Figs. 3B–C, S3, S4). Although metabolite titers were similar between the two conditions (ca. 8 mM acetone for N2 and ^13^CH_4_), ^13^C tracing revealed that methane-derived carbon was used to partially produce acetone. When using a ^13^CH_4_ atmosphere (Fig. 3D), the average carbon labeling in acetone was 9.1 ± 1.3%, which is significantly higher than the 1.9 ± 0.8% average carbon labeling when using a N_2_ atmosphere. The latter is similar to expected natural abundance (ca. 1.1%). As expected, ^13^C tracing revealed that the inactive sMMO control strain was unable to convert ^13^C-methane carbon to acetone (Fig. S5).

**Figure 3.**
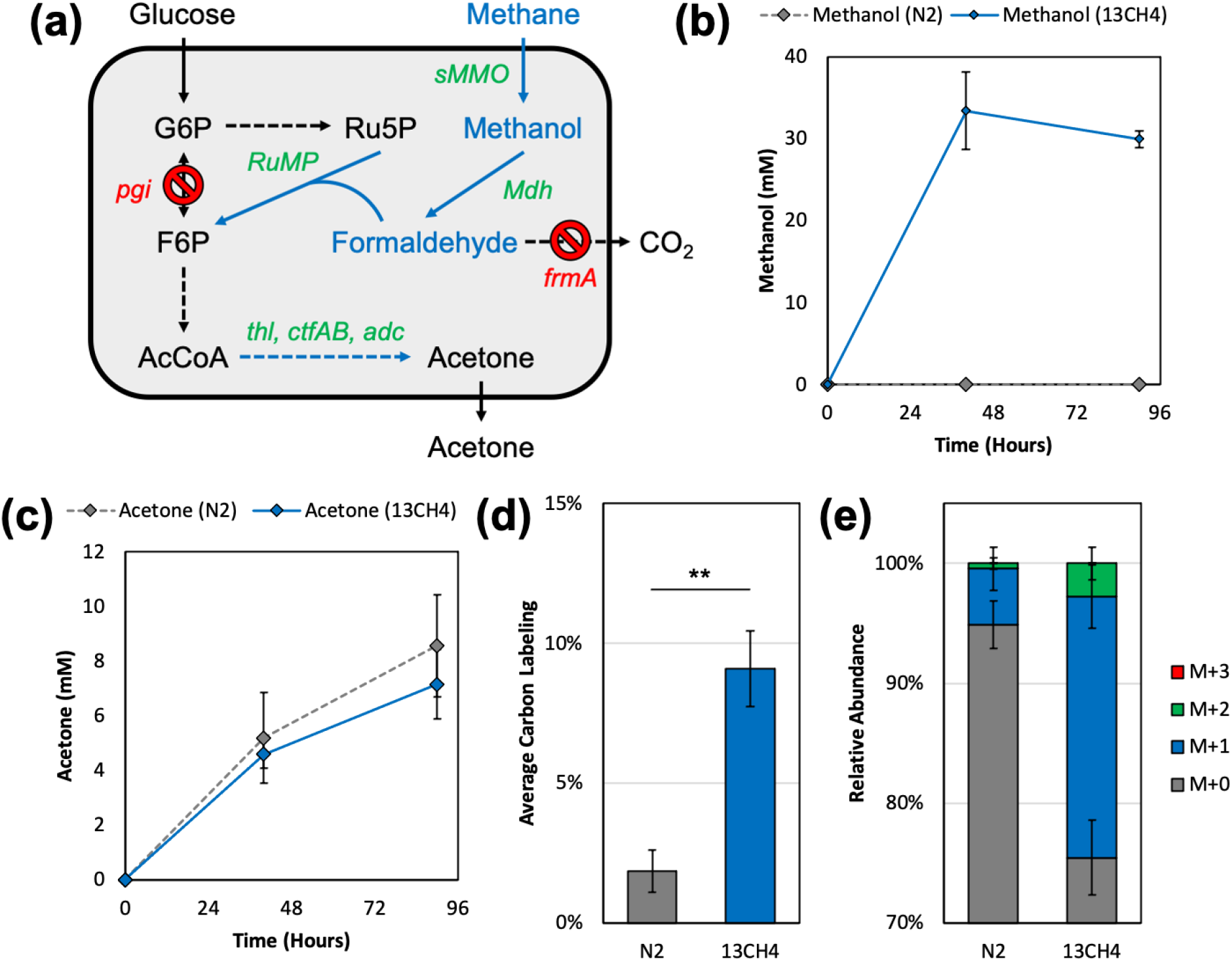
Methane bioconversion to acetone in a synthetic *E. coli* methanotroph. (A) Engineered *E. coli* co-utilizes methane and glucose for acetone production via deletion of phosphoglucose isomerase (*pgi*) and episomal expression of heterologous genes. Heterologous genes are shown in green with their corresponding pathways shown in blue. Production of methanol (B) and acetone (C) during gas fermentation. (D) Average ^13^C labeling of acetone and (E) relative abundance of acetone mass isotopomers at the end of the fermentation (90 hours). Abbreviations: sMMO (soluble methane monooxygenase), Mdh (methanol dehydrogenase), RuMP (ribulose monophosphate pathway consisting of hexulose phosphate synthase and phosphohexulose isomerase), thl (thiolase), ctfAB (CoA transferase), adc (acetoacetate decarboxylase). Error bars indicate standard deviation (n=5 for N_2_, n=10 for ^13^CH_4_). ** p < 0.01. See text for more details.

Since methane-dependence is not established with only Δ*pgi*, i.e., Δ*pgi* can still utilize glucose in the absence of methane, it is difficult to predict the expected ^13^C labeling pattern, though M+1 mass isotopomers should be most prevalent. However, the average carbon labeling in acetone was higher when using ^13^C-methane and glucose compared to that from ^13^C-methanol and glucose in a non-sMMO-expressing strain (2.4 ± 0.3%) ^25^, suggesting that *E. coli* may be better equipped for methane utilization. Furthermore, the relative abundance of acetone mass isotopomers reveals that ^13^C labeling is primarily M+1 (Figs. 3E, S5; 1-M0=25%), which is expected based on the co-utilization scheme that Δ*pgi* creates. Several engineering strategies can be used to increase the degree of methane conversion. One such strategy is to engineer a methane-dependent strain that co-utilizes methane and a sugar in a one-to-one molar ratio. Several methanol-dependent *E. coli* methylotrophs have been developed ^27–29^. Under these dependent conditions, the average ^13^C labeling in acetone increases to 33% ^27^. Alternatively, tuning the native or a heterologous non-oxidative PPP for increased cycling would generate higher order Ru5P mass isotopomers, which leads to improved methane utilization ^25,30^.

## Discussion

Ongoing efforts to improve the system towards commercially relevant MMO rates (400-500 mg methanol·gCDW^−1^·h^−1^) include expression balancing the GroESL chaperones, incorporating overexpression of additional protein folding chaperones, and additional rounds of directed evolution on the sMMO itself. The strategy used here could be applied to other difficult-to-express enzymes, including cytochromes P450. The sMMO has broad substrate specificity, making it an attractive catalyst for other biotechnology applications; the ability to engineer the substrate specificity is especially important here^31^. Finally, the system described can be used to understand other aspects of this fascinating enzyme, including testing mechanistic hypotheses for the catalytic cycle and participation of key residues. Ultimately, autonomous synthetic methanotrophy, i.e., the ability to utilize methane as the sole carbon and energy source, is desirable and may be realized via further engineering efforts or adaptive laboratory evolution (ALE). One such strategy is to equip a true *E. coli* methylotroph ^32^, achieved via ALE, with the sMMO. However, further engineering and evolutionary efforts will likely be required to obtain a robust *E. coli* methanotroph. Together, this study provides the first reported demonstration of *in vivo* methane bioconversion to liquid fuels and chemicals in a synthetic methanotroph and establishes the next step toward realizing industrial methane bioconversion.

## Acknowledgements

This work was supported by the Advanced Research Projects Agency-Energy (ARPA-E) Reducing Emissions using Methanotrophic Organisms for Transportation Energy (REMOTE) program (DE-AR0000432). This publication is based upon work supported by the Climate Change and Emissions Management (CCEMC) Corporation/Emissions Reductions Alberta under Project K130103 and the National Science Foundation under Grant No. 1520425. Gen9, Inc. (Ginkgo Bioworks) provided Materials for this work under the 2014 G-Prize. The authors would like to kindly thank Dr. Gwendolyn Gregory for assistance with GC-FID analysis.

## Author Contributions

RKB, ND, BZ, NH, DG, EC and ETP designed the research. RKB, ND, MD, SJ, KH, BZ, NH, DG and EC conducted the experiments. RKB, ND, EC and ETP analyzed the data and wrote the manuscript. All authors read and approved the manuscript.

## References

1 Haynes, C. A. & Gonzalez, R. Rethinking biological activation of methane and conversion to liquid fuels. Nat Chem Biol 10, 331–339, doi:10.1038/nchembio.1509 (2014).

2 Whitaker, W. B., Sandoval, N. R., Bennett, R. K., Fast, A. G. & Papoutsakis, E. T. Synthetic methylotrophy: engineering the production of biofuels and chemicals based on the biology of aerobic methanol utilization. Curr Opin Biotechnol 33, 165–175, doi:10.1016/j.copbio.2015.01.007 (2015).

3 Bennett, R. K., Steinberg, L. M., Chen, W. & Papoutsakis, E. T. Engineering the bioconversion of methane and methanol to fuels and chemicals in native and synthetic methylotrophs. Curr Opin Biotechnol 50, 81–93, doi:10.1016/j.copbio.2017.11.010 (2018).

4 Bennett, R. K. et al. Triggering the stringent response enhances synthetic methanol utilization in Escherichia coli. Metab Eng 61, 1–10, doi:10.1016/j.ymben.2020.04.007 (2020).

5 Woolston, B. M., King, J. R., Reiter, M., Van Hove, B. & Stephanopoulos, G. Improving formaldehyde consumption drives methanol assimilation in engineered E. coli. Nature Communications 9, 2387, doi:10.1038/s41467-018-04795-4 (2018).

6 Whitaker, W. B. et al. Engineering the biological conversion of methanol to specialty chemicals in Escherichia coli. Metab Eng 39, 49–59, doi:10.1016/j.ymben.2016.10.015 (2017).

7 Muller, J. E. N. et al. Engineering Escherichia coli for methanol conversion. Metab Eng 28, 190–201, doi:10.1016/j.ymben.2014.12.008 (2015).

8 Kim, H. J. et al. Biological conversion of methane to methanol through genetic reassembly of native catalytic domains. Nature Catalysis 2, 342–353, doi:10.1038/s41929-019-0255-1 (2019).

9 Banerjee, R., Jones, J. C. & Lipscomb, J. D. Soluble Methane Monooxygenase. Annu Rev Biochem 88, 409–431, doi:10.1146/annurev-biochem-013118-111529 (2019).

10 West, C. A., Salmond, G. P., Dalton, H. & Murrell, J. C. Functional expression in Escherichia coli of proteins B and C from soluble methane monooxygenase of Methylococcus capsulatus (Bath). J Gen Microbiol 138, 1301–1307, doi:10.1099/00221287-138-7-1301 (1992).

11 Murrell, J. C., Gilbert, B. & McDonald, I. R. Molecular biology and regulation of methane monooxygenase. Arch Microbiol 173, 325–332, doi:10.1007/s002030000158 (2000).

12 Kim, H. et al. MMOD-induced structural changes of hydroxylase in soluble methane monooxygenase. Sci Adv 5, eaax0059, doi:10.1126/sciadv.aax0059 (2019).

13 Merkx, M. & Lippard, S. J. Why OrfY? Characterization of MMOD, a long overlooked component of the soluble methane monooxygenase from Methylococcus capsulatus (Bath). J Biol Chem 277, 5858–5865, doi:10.1074/jbc.M107712200 (2002).

14 Lloyd, J. S., De Marco, P., Dalton, H. & Murrell, J. C. Heterologous expression of soluble methane monooxygenase genes in methanotrophs containing only particulate methane monooxygenase. Archives of Microbiology 171, 364–370, doi:10.1007/s002030050723 (1999).

15 Smith, T. J. & Murrell, J. C. in Methods in Enzymology: Methods in Methane Metabolism, Vol 495, Pt B Methods in Enzymology (eds A. C. Rosenzweig & S. W. Ragsdale) 135–147 (2011).

16 Smith, T. J., Slade, S. E., Burton, N. P., Murrell, J. C. & Dalton, H. Improved system for protein engineering of the hydroxylase component of soluble methane monooxygenase. Applied and Environmental Microbiology 68, 5265–5273, doi:10.1128/aem.68.11.5265-5273.2002 (2002).

17 Miller, A. R., Keener, W. K., Watwood, M. E. & Roberto, F. F. A rapid fluorescence-based assay for detecting soluble methane monooxygenase. Appl Microbiol Biotechnol 58, 183–188, doi:10.1007/s00253-001-0885-4 (2002).

18 Clarke, E. J., Zhu, B., Greenfield, D. L., Jones, S. R. & Helman, N. C. Functional Expression of Monooxygenases and Methods of Use. WO/2017/087731 (2017).

19 Erijman, A., Dantes, A., Bernheim, R., Shifman, J. M. & Peleg, Y. Transfer-PCR (TPCR): a highway for DNA cloning and protein engineering. J Struct Biol 175, 171–177, doi:10.1016/j.jsb.2011.04.005 (2011).

20 Clarke, E. J., Greenfield, D. L., Helman, N. C., Jones, S. R. & Zhu, B. Improved methane monooxygenase enzymes. WO/2019/010455A1 (2019).

21 Carlsen, H. N., Joergensen, L. & Degn, H. Inhibition by ammonia of methane utilization in Methylococcus capsulatus (Bath). Applied Microbiology and Biotechnology 35, 124–127, doi:10.1007/BF00180649 (1991).

22 Fox, B. G., Froland, W. A., Dege, J. E. & Lipscomb, J. D. Methane monooxygenase from Methylosinus trichosporium OB3b. Purification and properties of a three-component system with high specific activity from a type II methanotroph. J Biol Chem 264, 10023–10033 (1989).

23 Colby, J., Stirling, D. I. & Dalton, H. The soluble methane mono-oxygenase of Methylococcus capsulatus (Bath). Its ability to oxygenate n-alkanes, n-alkenes, ethers, and alicyclic, aromatic and heterocyclic compounds. Biochem J 165, 395–402, doi:10.1042/bj1650395 (1977).

24 Harwood, J. H. & Pirt, S. J. Quantitative Aspects of Growth of the Methane Oxidizing Bacterium Methylococcus capsulatus on Methane in Shake Flask and Continuous Chemostat Culture. Journal of Applied Bacteriology 35, 597–607, doi:10.1111/j.1365-2672.1972.tb03741.x (1972).

25 Bennett, R. K., Gonzalez, J. E., Whitaker, W. B., Antoniewicz, M. R. & Papoutsakis, E. T. Expression of heterologous non-oxidative pentose phosphate pathway from Bacillus methanolicus and phosphoglucose isomerase deletion improves methanol assimilation and metabolite production by a synthetic Escherichia coli methylotroph. Metab Eng 45, 75–85, doi:10.1016/j.ymben.2017.11.016 (2018).

26 Bermejo, L. L., Welker, N. E. & Papoutsakis, E. T. Expression of Clostridium acetobutylicum ATCC 824 genes in Escherichia coli for acetone production and acetate detoxification. Appl Environ Microbiol 64, 1079–1085 (1998).

27 Bennett, R. K. et al. Engineering Escherichia coli for methanol-dependent growth on glucose for metabolite production. Metab Eng 60, 45–55, doi:10.1016/j.ymben.2020.03.003 (2020).

28 Meyer, F. et al. Methanol-essential growth of Escherichia coli. Nat Commun 9, 1508, doi:10.1038/s41467-018-03937-y (2018).

29 Chen, C. T. et al. Synthetic methanol auxotrophy of Escherichia coli for methanol-dependent growth and production. Metab Eng 49, 257–266, doi:10.1016/j.ymben.2018.08.010 (2018).

30 Rohlhill, J., Gerald Har, J. R., Antoniewicz, M. R. & Papoutsakis, E. T. Improving synthetic methylotrophy via dynamic formaldehyde regulation of pentose phosphate pathway genes and redox perturbation. Metab Eng 57, 247–255, doi:10.1016/j.ymben.2019.12.006 (2020).

31 Smith, T. J. & Dalton, H. Vol. 151 Studies in Surface Science and Catalysis. (ed Rafael Vazquez-Duhalt & Rodolfo Quintero-Ramirez) 177–192 (Elsevier, 2004).

32 Chen, F. Y., Jung, H. W., Tsuei, C. Y. & Liao, J. C. Converting Escherichia coli to a Synthetic Methylotroph Growing Solely on Methanol. Cell 182, 933–946 e914, doi:10.1016/j.cell.2020.07.010 (2020).

