## Supplemental Information for "Expression of soluble methane monooxygenase in *Escherichia coli* enables methane conversion"

### Supplementary Figures

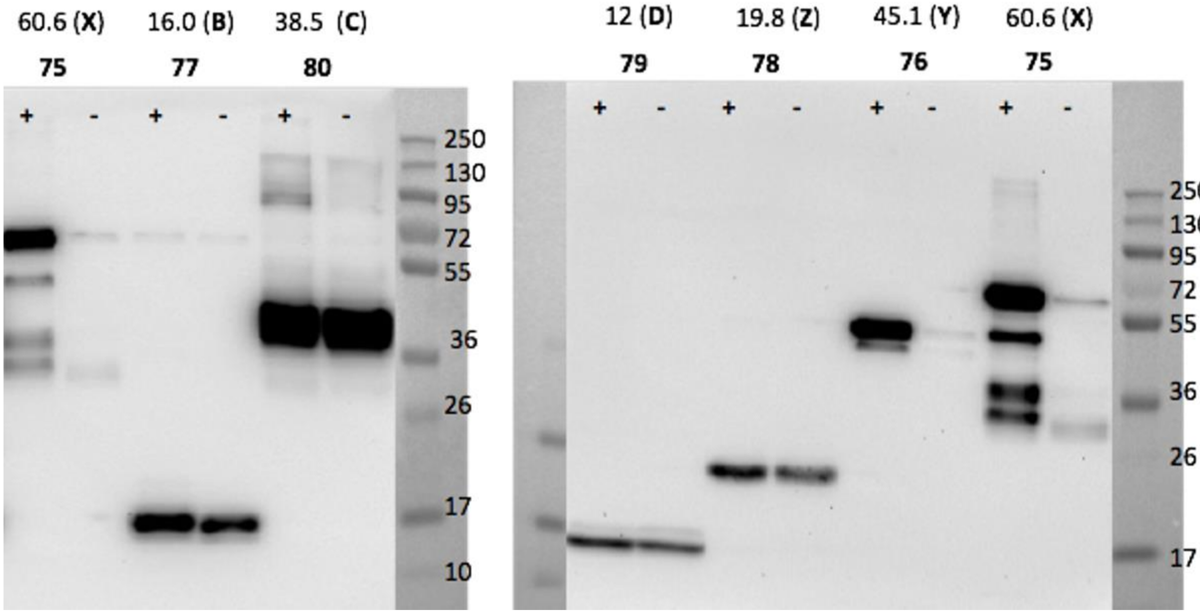


**Figure S1.** Western blotting of his-tagged sMMO components in the presence or absence of overexpressed chaperones. Boiled eluted fractions in 4× Laemmli buffer containing BME (20% v/v). Samples (5 uL) and PageRuler Plus prestained protein ladder (5 uL, Themo) were run on a 10-12% Bio-Rad Mini-Protean TGX stain-free precast gel in tris-glycine running buffer (250 mM Tris base, 1.92 M glycine, 1% SDS) at 150 V until ladder reached the bottom of the gel. The gel was equilibrated in transfer buffer (25 mM Tris, 192 mM glycine, 20% v/v methanol) for 20 min before transferring at 125 V for 1 h to a pre-wetted PVDF membrane (Bio-Rad) using a BioRad Mini Trans-Blot transfer cell. All subsequent steps utilized a rocking agitator to provide mixing. The membrane was blocked for 2 h at room temperature (5% w/v milk (Apex BioResearch Products, Research Triangle Park, NC) in TBST buffer (50 mM Tris, pH 8.0, 150 mM NaCl, 0.1% w/v Tween-20) followed by incubation with rabbit anti-6xHis primary antibodies (1:3333 dilution in TBST containing 5% w/v milk at room temperature for 3 h. The membrane was washed in TBST (3 × 10 min) before incubating with Bio-Rad Goat Anti-Rabbit IgG-HRP (Bio-Rad Laboratories,1:5000) at room temperature for 1.5 h. The membrane was washed again TBST (3 × 10 min) before visualization with a Western Lightning Plus ECL kit (PerkinElmer; Waltham, MA) using a Bio-Rad Mini Trans-Blot Cell and Quantity One software.

```
Score             Expect   Identities      Gaps         Strand
1349 bits(730)    0.0      736/739(99%)    0/739(0%)    Plus/Plus

Query  1    GATCAAAGGATCTTCTTGAGATCCtttttttCTGCGCGTAATCTTTTGCCCTGTAAACGA  60
            ||||||||||||||||||||||||||||||||||||||||||||||||||||||||||||
Sbjct  1    GATCAAAGGATCTTCTTGAGATCCTTTTTTTCTGCGCGTAATCTTTTGCCCTGTAAACGA  60

Query  61   AAAAACCACATTGGGAGGTGGTTTGATCGAAGGTTAAGTCAGTTGGGGAACTGCTTAACC  120
            ||||||||| | ||||||||||||||||||||||||||||||||||||||||||||||||
Sbjct  61   AAAAACCACCTGGGGAGGTGGTTTGATCGAAGGTTAAGTCAGTTGGGGAACTGCTTAACC  120

Query  121  GTGGTAACTGGCTTTCGCAGAGCACAGCAACCAAATCTGTCCTTCCAGTGTAGCCGGACT  180
            ||||||||||||||||||||||||||||||||||||||||||||||||||||||||||||
Sbjct  121  GTGGTAACTGGCTTTCGCAGAGCACAGCAACCAAATCTGTCCTTCCAGTGTAGCCGGACT  180

Query  181  TTGGCGCACACTTCAAGAGCAACCGCGTGTTTAGCTAAACAAATCCTCTGCGAACTCCCA  240
            ||||||||||||||||||||||||||||||||||||||||||||||||||||||||||||
Sbjct  181  TTGGCGCACACTTCAAGAGCAACCGCGTGTTTAGCTAAACAAATCCTCTGCGAACTCCCA  240

Query  241  GTTACCAATGGCTGCTGCCAGTGGCGTTTTACCGTGCTTTTCCGGGTTGGACTCAAGTGA  300
            ||||||||||||||||||||||||||||||||||||||||||||||||||||||||||||
Sbjct  241  GTTACCAATGGCTGCTGCCAGTGGCGTTTTACCGTGCTTTTCCGGGTTGGACTCAAGTGA  300

Query  301  ACAGTTACCGGATAAGGCGCAGCAGTCGGGCTGAACGGGGAGTTCTTGCTTACAGCCCAG  360
            ||||||||||||||||||||||||||||||||||||||||||||||||||||||||||||
Sbjct  301  ACAGTTACCGGATAAGGCGCAGCAGTCGGGCTGAACGGGGAGTTCTTGCTTACAGCCCAG  360

Query  361  CTTGGAGCGAACGACCTACACCGAGCCGAGATACCAGTGTGTGAGCTATGAGAAAGCGCC  420
            ||||||||||||||||||||||||||||||||||||||||||||||||||||||||||||
Sbjct  361  CTTGGAGCGAACGACCTACACCGAGCCGAGATACCAGTGTGTGAGCTATGAGAAAGCGCC  420

Query  421  ACACTTCCCGTAAGGGAGAAAGGCGGAACAGGTATCCGGTAAACGGCAGGGTCGGAACAG  480
            ||||||||||||||||||||||||||||||||||||||||||||||||||||||||||||
Sbjct  421  ACACTTCCCGTAAGGGAGAAAGGCGGAACAGGTATCCGGTAAACGGCAGGGTCGGAACAG  480

Query  481  GAGAGCGCAAGAGGGAGCGACCAGCCGGAAACGGTGGGGATCTTTAAGTCCTGTCGGGTT  540
            |||||||||||||||||||||| |||||||||||||||||||||||||||||||||||||
Sbjct  481  GAGAGCGCAAGAGGGAGCGACCCGCCGGAAACGGTGGGGATCTTTAAGTCCTGTCGGGTT  540

Query  541  TCGCCCGTACTGTCAGATTCATGGTTGAGCCTCACGGCTCCCACAGATGCACCGGAAAAG  600
            ||||||||||||||||||||||||||||||||||||||||||||||||||||||||||||
Sbjct  541  TCGCCCGTACTGTCAGATTCATGGTTGAGCCTCACGGCTCCCACAGATGCACCGGAAAAG  600

Query  601  CGTCTGTTTATGTGAACTCTGGCAGGAGGGCGGAGCCTATGGAAAAACGCCACCGGCGCG  660
            ||||||||||||||||||||||||||||||||||||||||||||||||||||||||||||
Sbjct  601  CGTCTGTTTATGTGAACTCTGGCAGGAGGGCGGAGCCTATGGAAAAACGCCACCGGCGCG  660

Query  661  GCCCTGCTGTTTTGCCTCACATGTTAGTCCCCTGCTTATCCACGGAATCTGTGGGTAACT  720
            ||||||||||||||||||||||||||||||||||||||||||||||||||||||||||||
Sbjct  661  GCCCTGCTGTTTTGCCTCACATGTTAGTCCCCTGCTTATCCACGGAATCTGTGGGTAACT  720

Query  721  TTGTATGTGTCCGCAGCGC  739
            |||||||||||||||||||
Sbjct  721  TTGTATGTGTCCGCAGCGC  739
```

**Figure S2.** Mutations in cloDF13 origin of pNH265 that increases copy number of the origin.

Top row is the WT cloDF13 origin, the lower row is the origin sequence in pNH265.

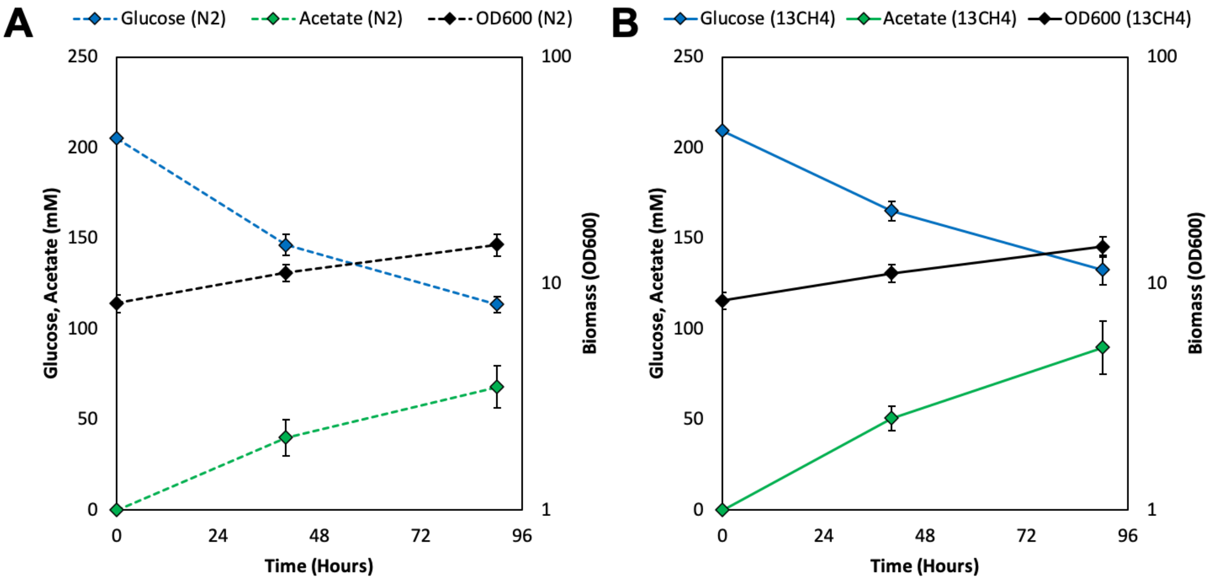


**Figure S3.** Fermentation profiles of Δ*frmA*Δ*pgi* + pUD11 + pNH284. Gas fermentation was performed in ammonium deficient M9 minimal medium supplemented with 1% yeast extract and 200 mM glucose. Following induction of the soluble methane monooxygenase, cells were concentrated and transferred to 160 mL serum bottles, which were then sealed with atmospheric headspace (to provide oxygen for methane oxidation) and pressurized with an additional 20 psi of either $N_2$ or $^{13}CH_4$. Cells were then grown at 37°C with shaking for 90 hours. Profiles of cell growth, glucose consumption, and acetate production during fermentation of cultures pressurized with $N_2$ (A) or $^{13}CH_4$ (B). Error bars indicate standard deviation (n=5 for $N_2$, n=10 for $^{13}CH_4$). See text for more details.

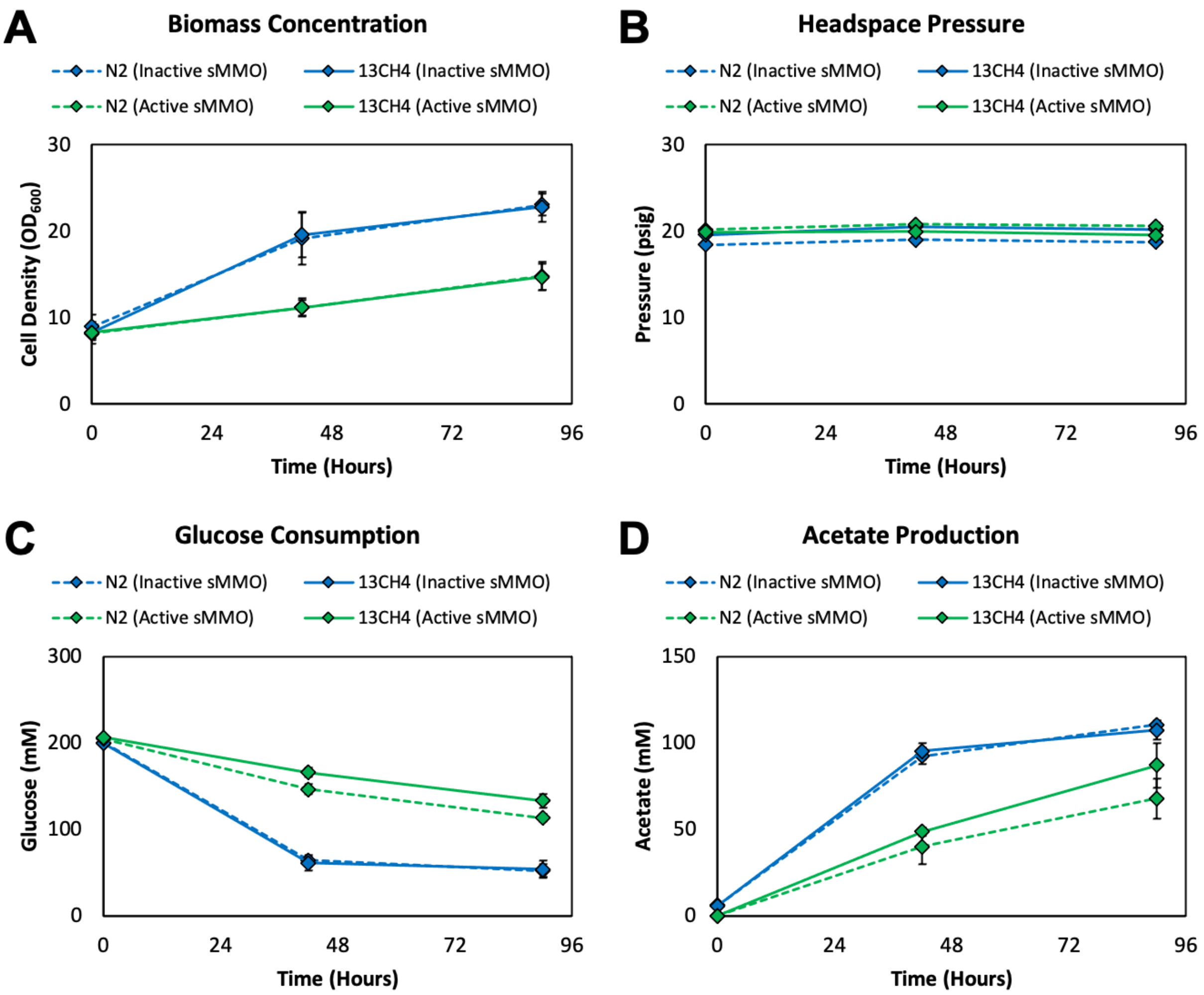


**Figure S4.** Fermentation profiles of $\Delta$*frmA*$\Delta$*pgi* + pUD11 + pNH284 ('Active sMMO') or + pLC215 ('Inactive sMMO'). Gas fermentation was performed in ammonium deficient M9 minimal medium supplemented with 1% yeast extract and 200 mM glucose. Following induction of the soluble methane monooxygenase, cells were concentrated and transferred to 160 mL serum bottles, which were then sealed with atmospheric headspace (to provide oxygen for methane oxidation) and pressurized with an additional 20 psi of either $N_2$ or $^{13}CH_4$. Cells were then grown at 37°C with shaking for 90 hours. Profiles of cell growth (A), headspace pressure (B), glucose consumption (C) and acetate production (D) during fermentation. Error bars indicate standard deviation (n=5 for $N_2$, n=10 for $^{13}CH_4$). See text for more details.

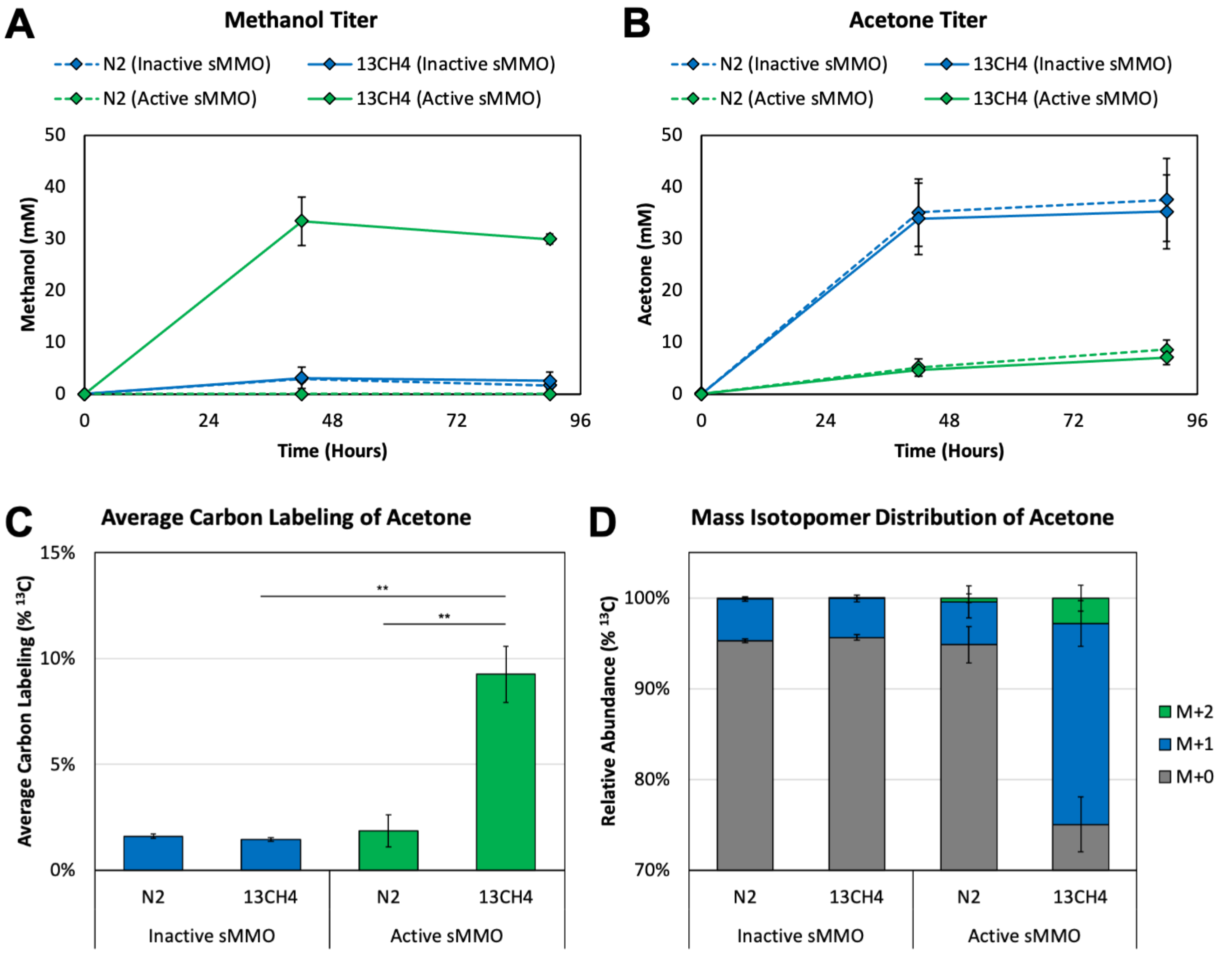


**Figure S5.** Fermentation profiles of Δ*frmA*Δ*pgi* + pUD11 + pNH284 (‘Active sMMO’) or + pLC215 (‘Inactive sMMO’). Gas fermentation was performed in ammonium deficient M9 minimal medium supplemented with 1% yeast extract and 200 mM glucose. Following induction of the soluble methane monooxygenase, cells were concentrated and transferred to 160 mL serum bottles, which were then sealed with atmospheric headspace (to provide oxygen for methane oxidation) and pressurized with an additional 20 psi of either $N_2$ or $^{13}CH_4$. Cells were then grown at 37°C with shaking for 90 hours. Profiles of methanol (A) and acetone (B) production during gas fermentation. (C) Average $^{13}C$ labeling of acetone and (D) relative abundance of acetone mass isotopomers at the end of the fermentation (90 hours). Error bars indicate standard deviation (n=5 for $N_2$, n=10 for $^{13}CH_4$). ** $p < 0.01$. See text for more details.

### Supplementary Tables

**Table S1.** Organisms from which monooxygenase and groESL genes were obtained.

| ***Organism*** | ***GroESL*** | ***Enzyme*** |
|---|---|---|
| *Methylococcus capsulatus (Bath)* | yes | MMO |
| *Methylopila trichosporium OB3b* | yes | MMO |
| *Pseudomonas mendocina* | none | Toluene MO |
| *Mycobacterium chubuense NBB4* | yes | Ethane MO |
| *Rhodococcus corallinus B-276* | none | Alkene MO |
| *Mycobacterium smegmatis MC*$^{2}$ *155* | yes | Propane MO |
| *Methylovulum miyakonense* | yes | MMO |
| *Methylomonas methanica MC09* | yes | MMO |
| *Methylocaldum sp175* | yes | MMO |
| *Methyloferula stellata* | yes | MMO |
| *Methylocystis sp LW5* | yes | MMO |
| *Solimonas aquatica DSM 25927* | yes | MMO |
| *Thauera butanivorans* | yes | Butane MO |
| *Methylocella silvestris BL2* | yes | Propane MO |
| *Acidomonas methanolica* | yes | Propane MO |
| *Methylocella silvestris BL2* | yes | MMO |
| *Xanthobacter autotrophicus* | yes | MMO |

**Table S2.** MMO activity in different base strains containing pNH284. Ratio of activity at 26 hours compared to NH848 is reported here. OverExpress was identified as the best strain, and methanol validation for this is shown in Fig. 2C of the main text.

| **Strain** | **Source** | **Genotype** | **Ratio of activity compared to NH848 (Coumarin assay)** |
|---|---|---|---|
| TOP10 | Invitrogen | F- mcrA Δ( mrr-hsdRMS-mcrBC) Φ80lacZΔM15 Δ lacX74 recA1 araD139 Δ( araleu)7697 galU galK rpsL (StrR) endA1 nupG | 0.05788241521 |
| ArcticExpress | Thermo Fisher Scientific | F– ompT hsdS(r – m –) dcm+ Tetr gal λ(DE3) endA Hte [cpn10 BB cpn60 Gentr] | 1.263248387 |
| OverExpress™ C43(DE3) Competent Cells | Biosearch Technologies | F – ompT hsdSB (rB- mB-) gal dcm (DE3) | 1.376727395 |
| Origami™ 2(DE3)pLysS Competent Cells | Sigma Aldrich | Δ(ara-leu)7697 ΔlacX74 ΔphoA PvuII phoR araD139 ahpC galE galK rpsL F′[lac+ lacIq pro] gor522::Tn10 trxB (StrR, TetR) | 0.1568851551 |
| Rosetta | Sigma Aldrich | F- ompT hsdSB(rB- mB-) gal dcm (DE3) pRARE2 (CamR) | 0.7417592129 |

**Table S3.** Engineering substrate specificity by altering MmoX. One mmoX mutation conferring ethane specificity was identified during site saturation mutagenesis; mmoX E240N. The mutant strain (BZ27) and wild type strain (BZ25) were assayed for ethane and methane oxidation using methods described above. Titers at 24 hours are shown.

| **Strain** | **mmoX mutation** | **Methanol (mM) / OD600** | **Ethanol (mM) / OD600** | **Specificity Ratio** |
|---|---|---|---|---|
| **BZ25** | Wild type | 5.45 | 0.94 | 17% |
| **BZ27** | E240N | 2.67 | 1.61 | 60% |

**Table S4.** Strains and plasmids used in this study.

| **Name** | **Relevant Characteristics** | **Reference** |
|---|---|---|
| **Strains** | | |
| Δ*frmA*Δ*pgi* | *E. coli* BW25113 Δ*frmA*::FRT Δ*pgi*::*kan* (Kan$^R$) | (*23*) |
| BZ36 | NH283/pDG6 | This study |
| BZ25 | NH283/pDG6 + pBZ13 | This study |
| BZ27 | NH283/pDG6-mmoX [E240N] + pBZ13 | This study |
| NH784 | NH283/pNH265 | This study |
| NH818 | NH283/pNH265-mmox [V23G] | This study |
| NH848 | NH283/pNH284 | This study |
| ND15 | OverExpress/pNH284 | This study |
| NH283 | NEB express DaraBAD::cat | This study |
| LC191 | NH283/pLC75 (pNH80 with 6x-His tagged mmoX) | This study |
| LC192 | NH283/pLC76 (pNH80 with 6x-His tagged mmoY) | This study |
| LC193 | NH283/pLC77 (pNH80 with 6x-His tagged mmoB) | This study |
| LC194 | NH283/pLC78 (pNH80 with 6x-His tagged mmoZ) | This study |
| LC195 | NH283/pLC79 (pNH80 with 6x-His tagged mmoD) | This study |
| LC196 | NH283/pLC80 (pNH80 with 6x-His tagged mmoc) | This study |
| **Plasmids** | | |
| pUD11 | pETM6_$P_{trc}$_*mdh_hps_phi*_$P_{thl}$\_*thl_ctfAB_adc* (ColE1, Amp$^R$) | (*23*) |
| pDG5 | Pbad_mmoXYBZDCgroEL2 (*M. capsulatus* Bath) (p15A, KanR) araC plus second CG origin repBL1 | This study |
| pDG6 | Pbad_mmoXYBZDC (*M. capsulatus* Bath) (p15A, KanR) araC plus second CG origin repBL1 | This study |
| pNH180 | PconstitutiveA_groESEL2 (*M. capsulatus* Bath) (cloDF13, specR) | This study |
| pBZ13 | PconstitutiveB_groESEL2 (*M. capsulatus* Bath) groESEL (E. coli) (cloDF13, specR) | This study |

| | | |
|---|---|---|
| pNH265 | Pbad_mmoXYBZDC (*M. capsulatus* Bath) PconstitutiveB_groESEL2 (*M. capsulatus* Bath)_groESEL (E. coli) araC (cloDF13, specR) | This study |
| pNH284 | pNH265 mmoX [V23G, T356G] mmoZ [R70E] mmoB [Y139S] groEL2 [N409G] | This study |
| pLC215 | pNH265 mmoX H246A - catalytically dead sMMO | This study |
| pNH157 | Pbad_mmoXYBZDC (*Methylocaldum* sp.175) (p15A, KanR) araC plus second CG origin repBL1 | This study |
| pNH158 | Pbad_mmoXYBZDC (*Methyloferula stellata*) (p15A, KanR) araC plus second CG origin repBL1 | This study |
| pNH80 | pDG6 with codon optimized mmoXYBZDC | This study |
| pNH185 | PconstitutiveA_groESEL2 (*Methylocaldum* sp.175) (cloDF13, specR) | This study |

**Table S5.** Promoter sequences for plasmids in Table S4.

| | |
|---|---|
| PconstitutiveA | TTGACAGCTAGCTCAGTCCTAGGGACTATGCTAGC (J23116) |
| PconstitutiveB | TTGACG |
